# Biological nitrification inhibition compromises the soil methane sink

**DOI:** 10.64898/2026.04.12.685468

**Authors:** Seongmin Yang, Fuhad H. Fahim, Pratap B. Shahi, Lauren E. Stanton, Suna Jo, Won-Min Park, Jesus A. Calleros, Sunghun Park, Jaejin Lee, Parnian Mohammadian, Prathap Parameswaran, Jaebeom Suh, Man Jae Kwon, Jeongdae Im

## Abstract

Biological nitrification inhibition (BNI) is a plant-mediated process that suppresses nitrification and is widely considered beneficial for reducing nitrous oxide emissions. Here, we show that BNI compounds also inhibit methane oxidation by methanotrophic bacteria, revealing a previously unrecognized trade-off in greenhouse gas regulation. Across soil bioreactor systems and pure cultures of both Type I and Type II methanotrophs, BNI compounds consistently suppressed methane oxidation activity. Kinetic analyses indicated an uncompetitive-like inhibition pattern, characterized by concurrent decreases in V_max_ and K_m_, while reversibility assays showed that inhibition was not associated with loss of cellular viability. Experiments under copper-replete and copper-depleted conditions further showed that inhibition is predominantly associated with particulate methane monooxygenase (pMMO). Transcriptomic analyses demonstrated compound-specific responses, including suppression of methane oxidation pathways and differential regulation of stress-associated genes. These findings suggest that BNI-mediated inhibition of methane oxidation may offset reductions in nitrous oxide emissions, with implications for predicting net greenhouse gas fluxes in agricultural and wetland ecosystems. Incorporating BNI effects into biogeochemical models will be critical for accurately evaluating their role in the global methane budget.

## Introduction

Plant–microbe interactions are fundamental drivers of terrestrial ecosystems, shaping nutrient cycles and influencing global climate dynamics. One critical and environmentally relevant interaction occurs in the plant rhizosphere, where plants and microbes compete for nitrogen (N). Despite heavy nitrogen fertilizer use in modern agriculture, approximately 50–70% of applied nitrogen is not utilized by crops, instead lost primarily due to nitrification^1,2^—a microbial process converting immobile ammonium (NH_4_^+^) to highly mobile nitrate (NO_3_^−^), contributing to nitrogen loss through leaching and denitrification. This transformation ultimately increases emissions of nitrous oxide (N_2_O), a potent greenhouse gas (GHG)^3^ and primary ozone depleting substance^4^.

To mitigate these losses, synthetic nitrification inhibitors (SNIs) have been developed; however, they face limitations due to limited biological stability, potential toxicity to beneficial soil organisms, and high costs^5,6^. In contrast, recent research has identified a natural plant-based alternative: Biological Nitrification Inhibition (BNI), where plants release specific root exudates capable of suppressing nitrifying microbes^6–9^. BNI represents a promising strategy to enhance nitrogen use efficiency and reduce N_2_O emissions^10–12^. For example, introducing chromosomal segments from wild wheat into high-yielding cultivars increased nitrogen uptake and reduced N_2_O emissions without compromising yield^8^.

While BNI shows promise in improving agricultural sustainability^2,13,14^, its broader ecological consequences remain insufficiently understood. Nitrification is initiated by the oxidation of NH_4_^+^ to NO_2_^−^ via NH_2_OH catalyzed by ammonia monooxygenase (AMO) and hydroxylamine oxidoreductase (HAO). Various BNI compounds have been identified with distinct inhibitory profiles^15,16^. For example, methyl 3-(4-hydroxyphenyl)propionate (MHPP) from sorghum (*Sorghum bicolor*) and 1,9-decanediol (1,9-D) from rice (*Oryza sativa*) primarily inhibit AMO^17,18^, whereas, linoleic acid (LA), derived from *Brachiaria humidicola*, inhibits both AMO and HAO^19^.

Methane, another major GHG, is oxidized by methanotrophs, which represent the primary biological methane sink and play a key role in mitigating emissions^20^. Methane oxidation is catalyzed by methane monooxygenases, including the membrane-bound particulate methane monooxygenase (pMMO) and the soluble methane monooxygenase (sMMO), whose expression is regulated by copper availability.^21^ Under typical environmental conditions, pMMO predominates and catalyzes the first step of methane oxidation. Given the chemical similarity between CH_4_ and NH_4_^+^, pMMO — which catalyzes the first step of methane oxidation—is evolutionarily related to AMO and shares structural and functional similarities.^22^ Accordingly, several SNIs—including nitrapyrin^23^, thiourea, allylthiourea^24^ and acetylene^25^—are known to inhibit methanotrophic activity as well. These parallels raise the possibility that naturally produced BNIs may also influence methane oxidation. Such interactions could have broad implications for terrestrial methane cycling, as methanotrophs are widely distributed across diverse soil and sediment environments and serve as the primary biological sink for methane emissions. Understanding such effects is particularly important in wetland and rice paddy systems^26^, where plants, methanotrophs, and ammonia oxidizers coexist and jointly regulate greenhouse gas fluxes.

To address this knowledge gap, we conducted a multi-scale investigation of BNI effects on methane oxidation. Using plant–soil systems, we first evaluated how plants with BNI activity influence methanotrophic activity and community structure. We then performed pure culture assays to quantify the inhibitory effects of representative BNI compounds on methanotrophs. To examine potential mechanisms, we performed molecular docking, simulations of BNI compounds, and analyzed their chemical analogs to identify interactions with pMMO. Finally, RNA-seq was used to characterize transcriptional responses to BNI compounds. Together, these results show that BNI compounds can suppress methane oxidation in addition to their established role in inhibiting nitrification, suggesting a previously underappreciated link between BNI activity and methane cycling. These findings highlight the importance of considering potential tradeoffs when evaluating BNI-driven impacts on coupled carbon and nitrogen processes in terrestrial ecosystems.

## Results

### Plants with BNI activity suppress methane oxidation in soil bioreactors

To determine whether plants with BNI activity influence methane oxidation in soil systems, we established soil bioreactors under three conditions: unamended control, NH_4_^+^ amendment (25 mg NH_4_^+^-N·kg_-soil_^−1^), and NH_4_^+^ amendment with *Brachiaria* (**Fig. 1a**). NH_4_^+^ addition significantly enhanced methane oxidation (P < 0.001), with the maximum methane consumption rate (V_max_) increasing from 18.0 (± 4.4) to 105.7 (± 8.8) nmol·min^−1^·g_-soil_^−1^. In contrast, *Brachiaria* incubation strongly suppressed methane oxidation (13.5 nmol·min^−1^·g_-soil_^−1^), reducing activity to levels below those observed in the unamended control, despite NH_4_^+^ amendment (P < 0.001) (**Fig. 1b**).

**Figure 1.**
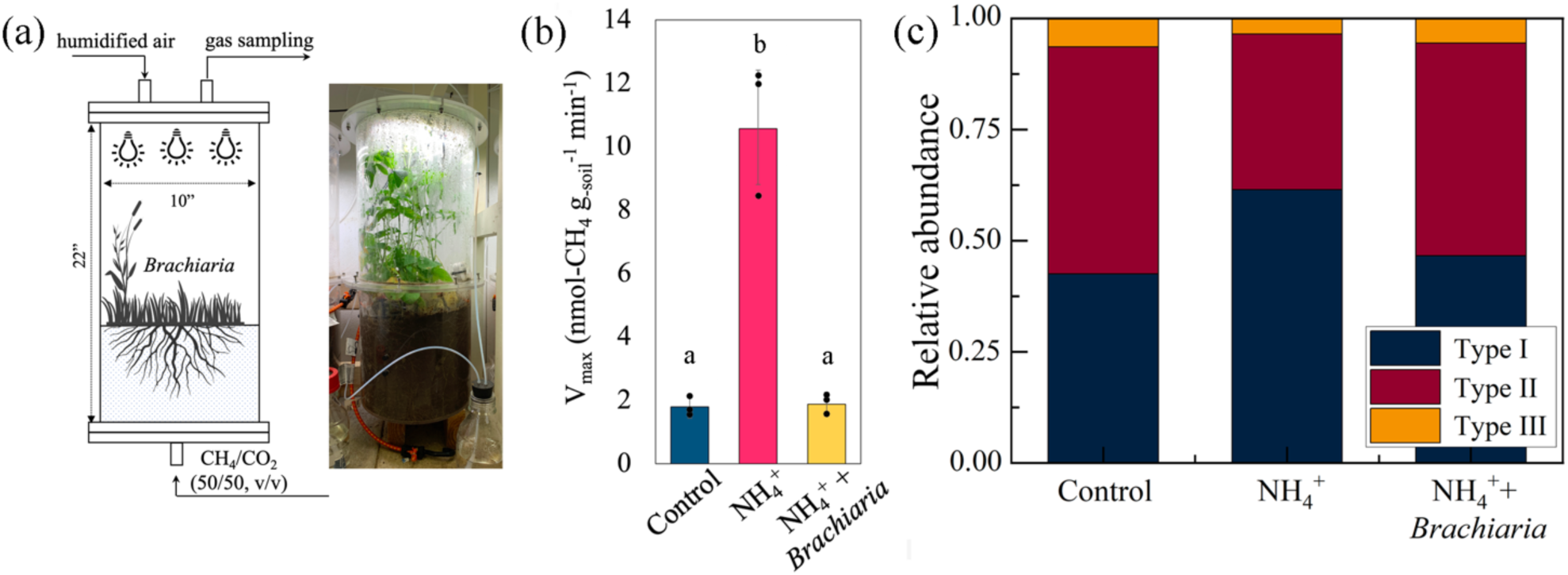
*Brachiaria* suppresses methane oxidation and alters methanotroph community structure in soil bioreactors. (a) Experimental design of soil bioreactors used to assess the impact of BNI on methane oxidation. Soils were incubated under three conditions: control, NH_4_^+^ amendment, and NH_4_^+^ amendment with the BNI-producing plant *Brachiaria*. (b) Maximum methane oxidation rates (V_max_) measured in soils recovered from the bioreactors. Bars indicate mean ± s.d. and dots represent individual biological replicates (n = 3). NH_4_^+^ amendment stimulated methane oxidation, whereas *Brachiaria* largely abolished this effect. Different letters indicate statistically significant differences (P < 0.05, one-way ANOVA with post hoc test). (c) Relative abundance of methanotroph lineages inferred from shotgun metagenomic sequencing based on extracted *pmoA* gene sequences. NH_4_^+^amendment increased the relative abundance of Type I methanotrophs, whereas incubation with *Brachiaria* reduced their contribution and increased the relative abundance of Type II lineages.

Shotgun metagenomic sequencing revealed pronounced treatment-dependent shifts in methanotroph composition (**Fig. 1c**). NH_4_^+^ addition alone led to a strong enrichment of Type I methanotrophs, consistent with previous observations that NH_4_^+^ availability stimulates fast-growing Type I lineages^27–29^. In contrast, soils incubated with NH_4_^+^ and *Brachiaria* showed a marked decline in Type I *pmoA* reads relative to NH_4_^+^-amended soils, while Type II methanotrophs were only moderately reduced. Type III group remained comparatively stable across treatments. Overall, the metagenomic patterns mirror the kinetic responses: NH_4_^+^ increased methane oxidation capacity, whereas *Brachiaria* suppressed it, with the suppression disproportionately affecting Type I methanotrophs in these soils. To determine whether the suppression of methane oxidation observed in soil resulted from direct inhibition of methanotrophs by BNI compounds, we next performed inhibition assays using axenic cultures of a representative methanotroph, *Methylosinus trichosporium* OB3b.

### BNI compounds inhibit methane oxidation in axenic methanotrophs

To evaluate the direct effects of BNI compounds on methanotrophic activity, we conducted inhibition assays using three structurally distinct compounds: MHPP, 1,9-D, and, LA, selected to represent different modes of inhibition previously reported in ammonia oxidizers^17–19^. Consistent with their distinct chemistries, the three BNI compounds exhibited distinct inhibition patterns in *M. trichosporium* OB3b. For all three compounds, increasing inhibitor concentration led to concurrent decreases in the maximum methane oxidation rate (V_max_) and apparent substrate affinity (K_m_), a kinetic signature consistent with an uncompetitive-like inhibition pattern, although the precise mechanism remains to be fully resolved (**Fig. 2a and 2b**)^30^. This kinetic behavior suggests that BNIs interact with pMMO at a site distinct from the methane-binding site, prompting structural analysis via molecular docking. Half-maximal inhibitory concentrations (IC_50_) were estimated to be approximately 730 μM for MHPP and 1070 μM for 1,9-D, based on concentration–response relationships derived from V_max_ inhibition and fitted using a four-parameter logistic (4PL) model (see **Supplementary Fig. S1, Supplementary Table S1,** and **Supplementary Methods S2**)^31^. In contrast, LA exhibited measurable inhibition at micromolar concentrations (**Fig. 2c**), indicating that LA interacts with OB3b at substantially lower concentrations than 1,9-D or MHPP. At its aqueous solubility limit (5.7 μM), LA reduced methane oxidation by ∼20%. As prior studies in *N. europaea* reported ∼80% inhibition at 55 µM^19^—above the aqueous solubility threshold tested here—we additionally tested LA at 57 µM for comparison. Under these conditions, inhibition increased to ∼42%, but did not reach 50%, precluding reliable IC_50_ estimation. Accordingly, LA was evaluated at its IC_40_, corresponding to 53 μM, in subsequent experiments. These data indicate that LA displays higher apparent potency but limited maximal inhibition under the experimental conditions tested.

**Figure 2.**
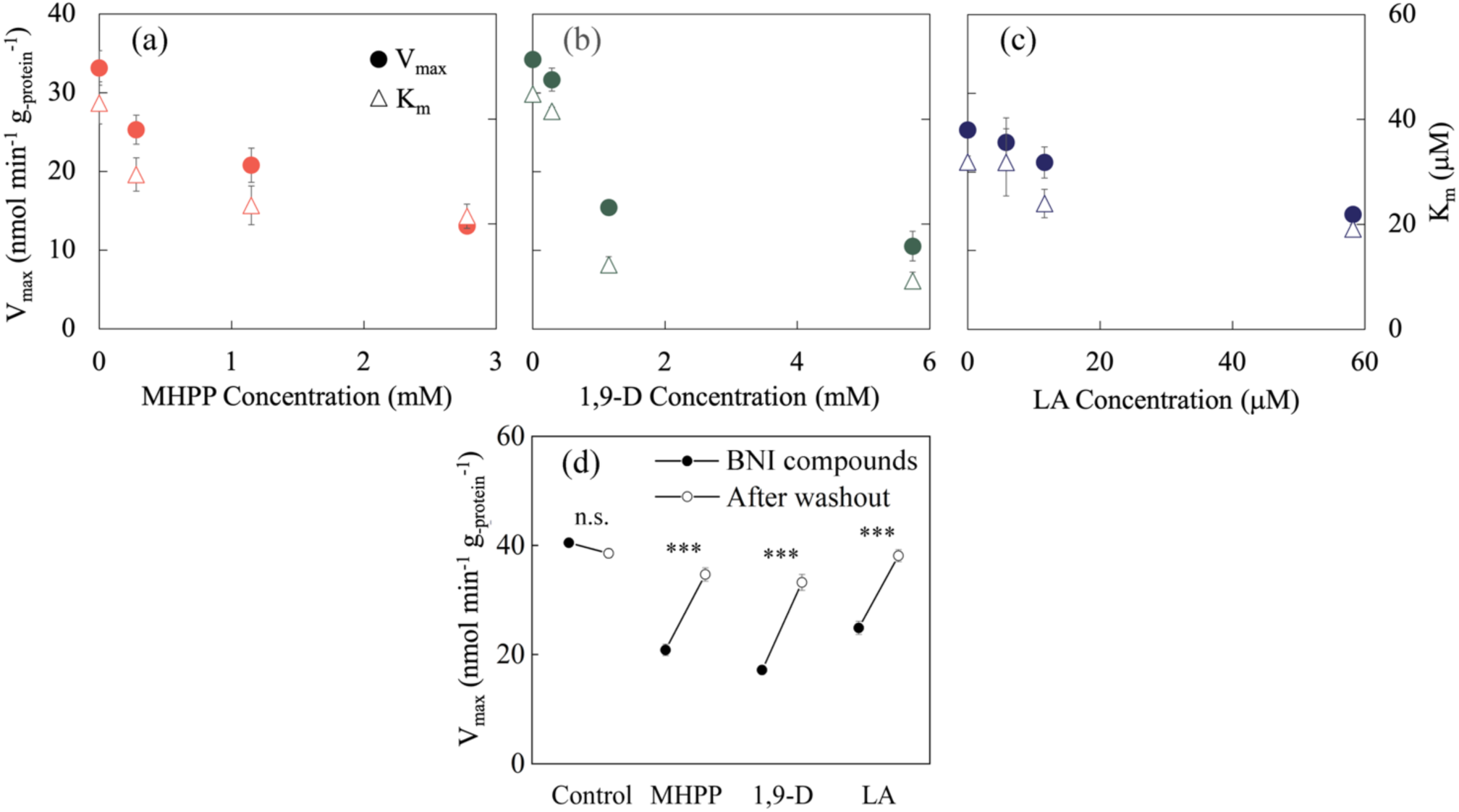
BNI compounds induce an uncompetitive-like inhibition pattern and reversible inhibition of methane oxidation. (a)–(c) Effects of individual BNI compounds on methane oxidation kinetics. Increasing concentrations of each compound resulted in concurrent decreases in V_max_ and K_m_, consistent with an uncompetitive-like inhibition pattern. (d) Reversibility of inhibition by BNI compounds following inhibitor removal. Methane oxidation activity recovered substantially after washout of each BNI compound, indicating that inhibition was reversible. Control samples showed no significant change following washout, whereas BNI-treated samples exhibited significant recovery of activity (paired two-tailed t-test, n = 3; *** P < 0.001; ns, not significant). Data are shown as V_max_ values. Error bars indicate s.d. and are shown where visible; in some cases they are smaller than the symbols.

To determine whether inhibition by BNI compounds was reversible or indicative of irreversible enzyme inactivation, OB3b cultures were incubated with each compound at their respective IC_50_ concentrations and to LA at its IC_40_ concentration, followed by removal of the compounds through washing and resuspension in fresh inhibitor-free medium prior to kinetic analysis. Following washout, methane oxidation activity recovered to 85–95% of untreated controls across treatments (**Fig. 2d**). These results indicate that BNI effects are reversible under the conditions tested and does not reflect suicidal inactivation of methane monooxygenase.

### BNI effects are predominantly associated with pMMO

While these results demonstrate that inhibition by BNI compounds is reversible, the enzymatic target remained unresolved. Given the structural and functional similarity between AMO and pMMO^22^, we reasoned that pMMO is the primary target of BNI effects. In contrast, the structurally distinct soluble methane monooxygenase (sMMO) would be less susceptible. To test this, *M. trichosporium* OB3b was assayed under copper-depleted conditions that repress pMMO and induce sMMO^21^. Cultures were exposed to MHPP and 1,9-D at their respective IC_50_ concentrations (730 and 1070 μM) and to LA at its IC_40_ concentration (53 μM). Activities were normalized to the corresponding untreated controls for each copper condition (**Fig. 3**). Under these conditions, methane oxidation activity in BNI-treated cultures was substantially higher under copper-depleted conditions relative to copper-replete conditions, indicating that inhibition was markedly reduced when pMMO activity was suppressed. The extent of recovery differed among compounds: MHPP-treated cultures retained near-control activity (88 ± 5%), whereas 1,9-D and LA exhibited lower relative activities (63 ± 9% and 79 ± 2%, respectively). These results indicate that the BNI effects are largely associated with pMMO activity, while 1,9-D and LA may exert additional inhibitory effects that persist under sMMO-dominant conditions.

**Figure 3.**
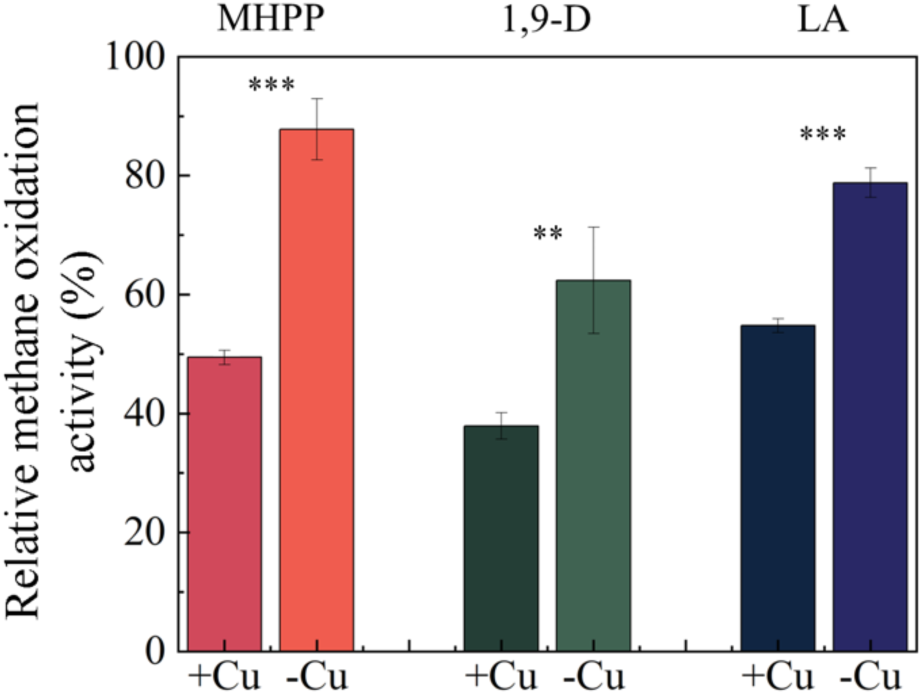
Inhibition by BNI compounds is reduced under sMMO-inducing conditions. Methane oxidation activity in *M. trichosporium* OB3b exposed to MHPP, 1,9-D, and LA under pMMO-active (+Cu) and sMMO-inducing (–Cu) conditions. Activities were normalized to the corresponding untreated controls for each copper condition. Inhibition by all compounds was reduced under sMMO-inducing conditions, consistent with a primary effect on pMMO. Error bars indicate s.d.. Asterisks indicate statistically significant differences between +Cu and −Cu conditions for each compound (**P < 0.01, ***P < 0.001; two-tailed unpaired t-test, n = 3).

### Molecular docking identifies a PmoB–PmoC interfacial binding pocket

To investigate the structural basis of the pMMO-associated inhibition observed in the methanotroph assays, molecular docking was performed using a homology model of *M. trichosporium* OB3b pMMO (**Fig. 4a**) with three representative BNI compounds: MHPP, 1,9-D, and LA. Across docking simulations, the highest-affinity poses were consistently located within the PmoB-PmoC interfacial pocket (**Fig. 4b and 4c**). Predicted binding energies for these compounds ranged from −4.6 to −6.1 kcal mol^−1^ (**Fig. 4d**).

**Figure 4.**
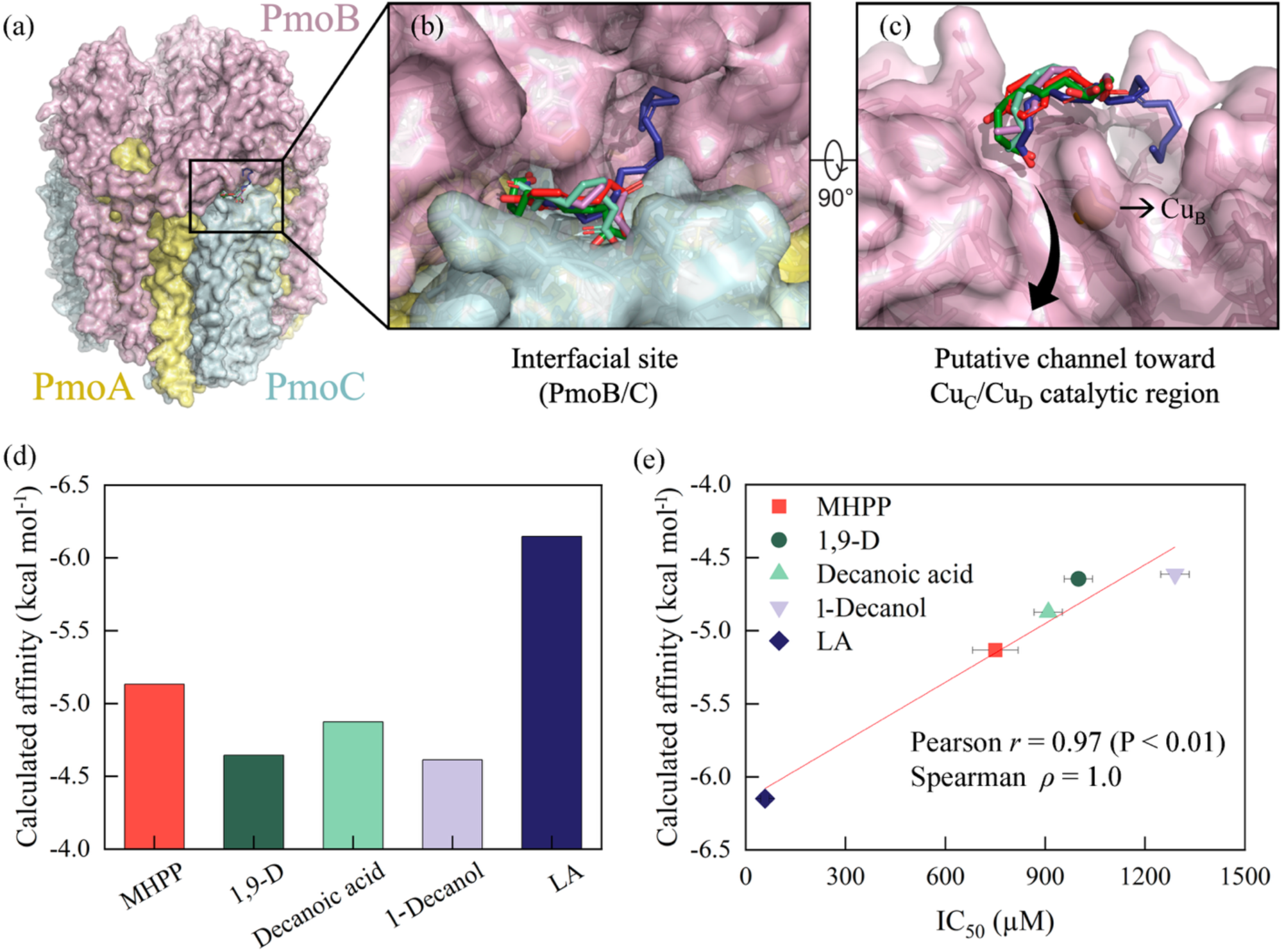
Molecular docking identifies a putative BNI-binding site in pMMO and links binding affinity to inhibitory potency. (a) Predicted homology model of pMMO showing the location of a putative binding site at the PmoB–PmoC interface. (b) Close-up view of docked poses for MHPP (red), 1,9-D (dark green), decanoic acid (light green), 1-decanol (light purple), and LA (navy blue) within the interfacial pocket. (c) Rotated view of the same pocket, showing its position near a putative channel toward the Cu*_C_*/Cu*_D_* catalytic region and in proximity to the Cu*_B_* site. (d) Calculated binding affinities of the docked compounds. (e) Relationship between calculated binding affinity (kcal mol^−1^) and inhibitory potency (IC_50_, except LA where IC_40_ is shown). Stronger binding was associated with lower IC_50_ values. Error bars indicate s.d. and are shown where visible; in some cases they are smaller than the symbols. Colors are consistent across panels.

To test whether this predicted binding site reflects a broader structural feature of hydrophobic inhibitors, two structural analogues of 1,9-D—dodecanoic acid and 1-decanol—were subsequently examined. Docking simulations predicted that both analogues bind within the same PmoB–PmoC interfacial pocket. Consistent with this prediction, both compounds inhibited methane oxidation in *M. trichosporium* OB3b (IC_50_ values of 910 μM and 1290 μM for dodecanoic acid and 1-decanol, respectively). These results support the predicted interaction of hydrophobic inhibitors with the PmoB–PmoC interfacial pocket.

Including the two structural analogues, predicted binding energies for all five compounds were strongly correlated with experimentally determined IC_50_ values (**Fig. 4e**), indicating that both the native BNI compounds and their structural analogues follow the predicted structure–activity relationship. Binding energy exhibited a perfect monotonic relationship with IC_50_ (Spearman’s *ρ* = 1.00, two-tailed P = 0.017) and a strong linear correlation between binding energy and IC_50_ (Pearson’s *r* = 0.97, P = 0.006). Although docking energies provide relative estimates rather than direct measures of binding affinity, the observed correlation supports the functional relevance of the PmoB–PmoC interfacial pocket as a plausible interaction site for BNI compounds.

The identified interfacial pocket lies adjacent to a channel-like cavity extending toward the Cu*_C_*/Cu*_D_* region of the PmoC subunit, proposed to harbor the catalytic copper center of pMMO^32,33^, and is also located near the Cu*_B_* site (**Fig. 4c**)^34^. Ligand binding at this site may influence molecular flux through this channel, for example by affecting substrate access or product egress, without directly occupying the catalytic center. This spatial arrangement is consistent with the inhibition pattern observed in the kinetic assays.

### Exposure to BNI compounds triggers compound-specific transcriptional responses

To further examine transcriptional responses of methanotrophs to BNI compounds, RNA-seq analysis was performed in biological triplicates. Transcriptomic profiling was conducted under conditions corresponding to the IC_50_ of MHPP and 1,9-D and the IC_40_ of LA, at which methane oxidation was inhibited by 47.5%, 58.6%, and 38.6%, respectively. RNA samples were collected at a time point corresponding to a period of high methane oxidation activity (10 h), as determined from preliminary time-course experiments (see **Supplementary Fig. S2**). Principal component analysis (PCA) revealed clear separation of MHPP-treated samples from controls, whereas 1,9-D induced broader dispersion consistent with widespread transcriptional perturbation. LA-treated samples showed a more moderate shift in global transcriptomic space (**Fig. 5a**).

**Figure 5.**
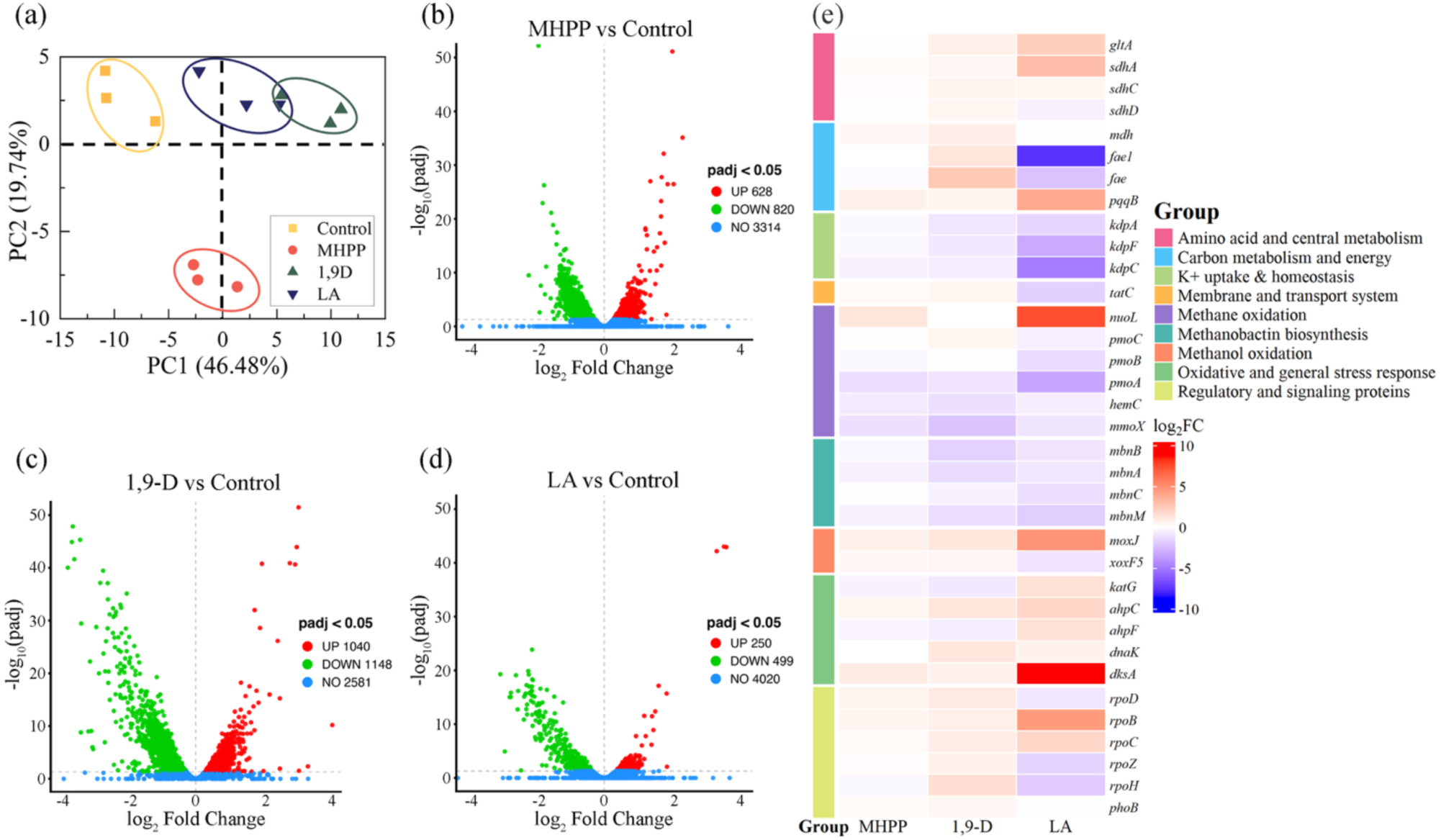
Compound-specific transcriptomic responses of *Methylosinus trichosporium* OB3b to BNI compound exposure. (a) Principal component analysis (PCA) showing global transcriptomic differences among treatments, with PC1 and PC2 explaining **XX%** and **XX%** of the total variance, respectively. (b–d) Volcano plots showing differential gene expression relative to control (adjusted *P* < 0.05). (e) Heatmap of selected genes involved in methane oxidation, central metabolism, transport, and stress response. Notable responses include upregulation of *nuoL* and *dksA* and downregulation of *pmoA* and *kdp* operon genes under LA treatment.

Differential expression analysis further highlighted these differences (**Fig. 5b-d**). A total of 1,448, 2,188, and 749 genes were significantly altered under MHPP, 1,9-D, and LA treatments, respectively, with 1,9-D eliciting the most extensive transcriptional response. Consistent with this, 1,9-D affected a large number of genes with moderate fold changes across diverse functional categories, including stress responses, transporters, and membrane-associated processes. MHPP induced a more structured transcriptional shift, with coordinated changes in specific metabolic pathways. In contrast, LA altered fewer genes overall but produced high-amplitude and high-significance expression changes in a subset of loci.

These patterns were further resolved at the gene level (**Fig. 5e**), where LA induced pronounced upregulation of *nuoL* and *dksA* and strong downregulation of *pmoA* and *kdp* operon genes, consistent with a selective but high-impact transcriptional response. Under these conditions, methane oxidation was inhibited by 48.5%, 58.6%, and 38.6% for MHPP, 1,9-D, and LA, respectively. Taken together, these results indicate that exposure to BNI compounds is associated with fundamentally distinct transcriptional programs in methanotrophs: 1,9-D acting as a broad stressor, MHPP producing a structured metabolic shift, and LA eliciting a selective but high-amplitude response.

### Type I methanotroph BG8 exhibits enhanced sensitivity to BNI compounds

Because metagenomic analysis of the soil bioreactors indicated that Type I methanotrophs were disproportionately reduced under *Brachiaria* incubation, inhibition assays were performed with *Methylomicrobium album* BG8, a representative Type I methanotroph, to determine whether this lineage-level pattern could be reproduced in pure culture. Across all compounds tested, BG8 was more sensitive to BNI compounds than *M. trichosporium* OB3b (Type II). The IC_50_ for MHPP was lower in BG8 (432 μM) than in OB3b (730 μM). Similarly, the IC_50_ for 1,9-D was lower in BG8 (391 μM) than in OB3b (1076 μM). For LA, the IC_40_ was also lower in BG8 (14 μM) than in OB3b (53 μM). These values were further normalized to cell abundance for cross-species comparison, as discussed below. Although these results are based on representative strains, this pattern is consistent with the preferential reduction of Type I taxa observed in soil metagenomes under *Brachiaria* incubation. Together, these results indicate that the tested methanotrophs are sensitive to BNI compounds. To place this response in context, we compared the inhibitory effects of BNIs on methanotrophs and ammonia-oxidizing bacteria, the canonical targets of these compounds.

### BNI compounds suppress methane oxidizers more strongly than ammonia oxidizers

To directly compare the inhibitory effects of BNI compounds on ammonia-oxidizing bacteria and methanotrophs, parallel inhibition assays were performed with *N. europaea*, a model ammonia-oxidizing bacterium, using the same three BNI compounds evaluated with methanotrophs: MHPP, 1,9-D, and LA. Because inhibitory responses depend on the ratio of inhibitor concentration to microbial biomass^35,36^, inhibitory potency was normalized to cell abundance, estimated from16S rRNA copy numbers, prior to comparison. In these assays, *N. europaea* cultures were tested at OD_600_ = 0.1, whereas methanotroph assays were performed at higher biomass (OD_600_ = 0.3).

For MHPP and 1,9-D, inhibitory potency was expressed as the negative log_10_ of the cell-normalized IC_50_ values (−log_10_ IC_50_) to facilitate comparison across orders of magnitude. Under this normalization, both methanotrophic strains were markedly more sensitive than *N. europaea*, indicating that BNI compounds suppress methane oxidation at lower per-cell exposure levels than those required to suppress nitrification (**Fig. 6a**).

**Figure 6.**
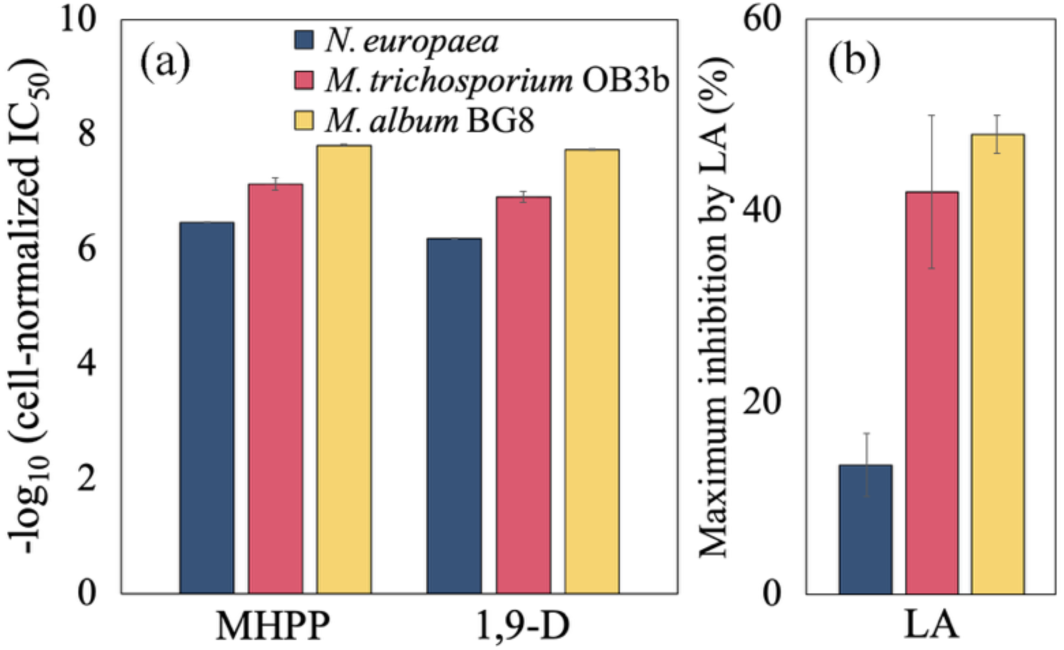
Comparative sensitivity of ammonia- and methane-oxidizing bacteria to BNI compounds. (a) Cell-normalized inhibitory potency of MHPP and 1,9-D toward *Nitrosomonas europaea*, *Methylosinus trichosporium* OB3b, and *Methylomicrobium album* BG8. Inhibitory potency is expressed as −log_10_(IC_50_) of the cell-normalized values (μM cell^−1^). Error bars represent propagated s.d. following log transformation of IC_50_ values and are shown where visible; in some cases they are smaller than the symbols. (b) Maximum inhibition by LA observed within the tested concentration range. Error bars represent s.d. of triplicate measurements. Error bars represent s.d. and may not be visible when smaller than the plotted symbols.

For LA, inhibition did not reach 50% in either ammonia-oxidizing bacteria or methanotrophs assays, precluding IC_50_ estimation. In *N. europaea*, inhibition reached its maximum (∼13%) at 57 μM, corresponding to a biomass-normalized exposure of 3.6 × 10^−8^ μM cell^−1^. Although previous studies using a bioluminescent, genetically modified *N. europaea* strain reported substantially higher inhibition (∼80%) at similar concentrations (55 μM)^19^, the native strain used here exhibited considerably lower sensitivity, likely reflecting differences in assay systems and strain backgrounds. Achieving the same normalized exposure in methanotrophs assays as in the *N. europaea* experiment would require substantially higher LA concentrations, estimated at approximately 734 μM for OB3b and 985 μM for BG8. However, inhibition in both methanotrophic strains had already reached a plateau within the tested concentration range (approximately 42% for OB3b and 48% for BG8), with maximal suppression observed at ∼57 μM (**Fig. 6b**). Thus, even at normalized exposures far below those required to match the ammonia-oxidizing bacteria assay, methanotrophs exhibited stronger inhibition than *N. europaea*. Together, these results suggest that BNI compounds may suppress methanotrophic activity to a similar or greater extent than nitrification in these assays.

## Discussion

This study shows that BNI compounds released by plants suppress not only ammonia oxidation but also methane oxidation by methanotrophs. By integrating soil bioreactor experiments, metagenomics, pure-culture kinetics, and molecular docking, we show that exposure to BNI compounds reduces the microbial capacity for methane oxidation. These findings reveal a previously unrecognized ecological tradeoff: while BNIs can mitigate nitrification and associated N_2_O emissions^12,37–39^, they may also diminish methane consumption. Because nitrification and methanotrophy are key microbial processes linking the nitrogen and carbon cycles, this dual effect has implications for greenhouse gas dynamics beyond local soil environments.

Our comparative assays extend this tradeoff beyond nitrification. Although BNIs were originally characterized as inhibitors of AMO in ammonia-oxidizing bacteria, methanotrophs exhibited greater apparent sensitivity on a per-cell basis across the compounds tested (**Fig. 6**). Cell-normalized inhibitory potencies differed by up to orders of magnitude between ammonia- and methane-oxidizing organisms under the conditions examined. Conceptually, the use of cell-normalized inhibitory potency parallels approaches widely used in pharmacology to compare compound efficacy across systems with differing biomass or target abundance^35,36^. However, such normalization is less commonly applied in environmental microbiology, where process rates are typically evaluated at the community or bulk level. As a result, while cell-based estimates provide a useful comparative framework, they should be interpreted cautiously when extrapolating to organismal or ecosystem-level responses. Rather, these results highlight the potential for BNI compounds to influence methane oxidation alongside nitrification, while underscoring the need for more comprehensive ecological evaluation.

The concentration ranges of BNI compounds tested in this study were selected to capture inhibitory responses spanning partial to near-maximal suppression, enabling quantitative comparison across compounds and organisms. Although *in situ* concentrations of individual BNI compounds in soils remain poorly constrained, previous soil-based assays provide a useful functional benchmark. The highest levels of biological nitrification inhibition have been reported in specialized tropical pasture grasses, particularly *B. humidicola*, which can release substantial amounts of nitrification inhibitors (e.g., 17–50 ATU g^−1^ root d^−1^) and achieve near-complete suppression (> 90%) of nitrification in soil systems^9^. In contrast, more moderate inhibition has been observed in crop species such as wheat, which produces MHPP and has been associated with 24–77% suppression of nitrification in soil assays^8,40^. These observations suggest that the concentration ranges used here fall within a functionally relevant range for comparative assessment of inhibitory potency, even if the contribution of individual compounds under field conditions remains unresolved. Importantly, BNI in soils likely reflects the combined effects of multiple compounds and their interactions, rather than the action of any single molecule, and thus the concentrations tested here should be interpreted as representative of effective inhibitory exposure rather than direct proxies for *in situ* concentrations.

Building on these observations, the combined kinetic and structural analyses provide a coherent framework for understanding inhibition of methane oxidation by BNI compounds. The uncompetitive-like inhibition pattern observed in kinetic assays (**Fig. 2**), together with the consistent localization of multiple compounds within the PmoB–PmoC interfacial pocket (**Fig. 4b**), suggest that BNI compounds act through peripheral interactions with pMMO that constrain catalytic turnover rather than directly competing with methane. The convergence of structurally distinct compounds on this region, along with the strong correlation between predicted binding energy and inhibitory potency, further supports the functional relevance of this interaction site. The position of this interfacial pocket adjacent to a channel leading toward the catalytic region raises the possibility that BNI binding may perturb molecular flux through this channel including effects on substrate access dynamics or product egress^41^, rather than directly occluding the active site. This region is also in close proximity to the Cu*_B_* site (**Fig. 4c**), a conserved mononuclear copper center coordinated by histidine residues and implicated in electron transfer and oxygen activation^34^. This spatial arrangement suggests that binding of BNI compounds at this site could interfere not only with substrate or product movement but also with electron transfer processes associated with catalytic turnover. Given that pMMO activity depends on coordinated electron delivery through copper centers, including Cu*_B_* and additional sites within the PmoC subunit, perturbation of this interfacial region may propagate through the copper network and reduce catalytic efficiency. Consistent with this interpretation, structural studies of the homologous AMO system have revealed coupling between copper centers through proton transfer pathways^42^, suggesting that coordinated transport and redox processes are a general feature of membrane-bound monooxygenases. However, because these inferences are based on indirect structural predictions and kinetic signatures, experimental structural validation of BNI compound–pMMO interactions will be required to fully establish the mechanism.

Despite this overall pattern, compound-specific differences were evident. MHPP acted in a largely pMMO-specific manner, whereas 1,9-D exhibited additional nonspecific effects, including residual inhibition under sMMO-inducing conditions (**Fig. 3**) and weaker predicted binding affinity (**Fig. 4d**). Notably, 1,9-D has previously been described as an AMO-specific inhibitor in ammonia-oxidizing bacteria^17,43^, suggesting a more restricted target range. The broader inhibitory effects observed here in methanotrophs indicate that its mode of action may be more context-dependent than previously recognized. In contrast, LA showed limited maximal inhibition, likely constrained by solubility, yet displayed high potency at low concentrations. In ammonia-oxidizing bacteria, LA has been reported to produce substantially higher levels of inhibition up to 80% at comparable concentrations (55 μM)^19^, suggesting that its inhibitory effect is strongly dependent on the biological system and experimental context. Transcriptomic responses further supported these distinctions: 1,9-D elicited broad stress-associated responses, MHPP produced a more targeted shift consistent with pMMO inhibition, and LA induced selective but high-amplitude response. Together, these results suggest that BNI compounds comprise a chemically diverse class of inhibitors with distinct modes of interaction and inhibitory profiles, rather than a single uniform mechanism.

At larger scales, these findings have implications for methane cycling in ecosystems where BNI activity is prevalent or strongly expressed. Wetlands, the largest natural source of atmospheric methane, and rice paddies, the largest anthropogenic source, both rely on methanotrophy in surface soils and rhizospheres as a key sink that moderates gross methane production^44,45^. Our results raise the possibility that BNI compounds released from plants may interfere with this sink function, thereby influencing net methane fluxes. In wetlands, BNI compound exudation by native vegetation could affect not only nitrogen cycling but also methanotrophy, potentially altering the balance of greenhouse gas fluxes. Accordingly, models of methane dynamics in wetlands and flooded systems may need to consider BNI-driven constraints on methanotrophy to more accurately capture net fluxes. In rice paddies, the implications are more direct, as rice itself produces BNI compounds such as 1,9-D^17,43^. Thus, BNI-based breeding strategies aimed at improving nitrogen-use efficiency and reducing N_2_O emissions may also influence methane oxidation, highlighting the importance of evaluating potential tradeoffs between nitrogen management and methane sink strength. Together, these considerations suggest that BNI activity could represent an underrecognized factor influencing methane–climate feedbacks.

Finally, several limitations of this study also point toward promising avenues for future research. The bioreactor and pure-culture experiments, although mechanistically informative, may not fully capture the complexity of plant–soil–microbe interactions in natural environments, where fluctuating oxygen availability, soil redox dynamics, microbial competition, and spatial heterogeneity in root exudation could all influence the magnitude of BNI effects. While our molecular docking analyses support the kinetic results consistent with an uncompetitive-like inhibition mechanism, they rely on predicted structures and simplified binding assumptions, and experimental structural confirmation of BNI compound–enzyme interactions will be an important next step. Moving forward, extending these findings to field-scale systems will be essential, combining methane flux measurements with high-resolution microbial and transcriptomic analyses in BNI-active environments such as rice paddies and wetlands. Incorporating BNI effects into process-based and Earth-system models will also be critical for evaluating their contribution to the global methane budget. As BNI-based strategies continue to be developed for improving nitrogen-use efficiency and reducing N_2_O emissions, interdisciplinary studies linking agronomy, microbiology, and climate science offer a promising path to ensure that their broader impacts on GHG dynamics are fully understood.

## Materials and Methods

### Soil bioreactor experiments

Acrylic columns (10 in internal diameter, 22 in length; **Fig. 1a**) were packed to a height of 25 cm with soil (bulk density 1.9-2.0 g cm^−3^) over a 2 cm glass bead layer to facilitate gas distribution. Columns were continuously supplied with a CH_4_/CO_2_ mixture (50:50, v/v) at 5 ± 0.5 ml min^−1^, while the headspace was flushed with humidified air at 20 ml·min^−1^. Soil moisture was maintained at 45–55% water holding capacity. Treatments (n=3) included (i) unamended control, (ii) NH_4_^+^ amendment (25 mg NH_4_^+^-N (kg_-soil_)^−1^), and NH_4_^+^ amendment with *Brachiaria* seedlings. After two months of incubation, soil samples were collected from the upper 5 cm, homogenized, and subsampled for downstream analyses: 10 g was used for methane oxidation kinetics and 5 g was used for metagenomic analysis. Detailed soil characterization, reactor operation, and sampling procedures are provided in **Supplementary Methods S1**.

### Metagenomic sequencing and community analysis

Metagenomic sequencing was performed to characterize methanotroph community composition in soil bioreactor samples. Genomic DNA was extracted using the MagMAX™ Microbiome Ultra Nucleic Acid Isolation Kit (Applied Biosystems, USA) with the KingFisherTM Duo Prime system (Thermo Fisher Scientific, USA). Sequencing libraries were prepared and sequenced on an Illumina platform (Novogene Co., Ltd., China). Paired-end reads were merged using FLASH, and merged FASTQ files were converted to FASTA format using Seqtk. High-quality sequences were used for downstream analysis. Custom *pmoA* reference databases were constructed from curated methanotroph sequences (see **Supplementary Table S2**) and expanded using FunGene-derived sequences to improve taxonomic resolution. Metagenomic sequences were aligned against the custom *pmoA* databases using BLASTN (e-value < 1 × 10^−6^ ; identity > 80%), and the highest-scoring hit was retained for classification. Relative abundances of methanotroph lineages were compared across treatments. Detailed sequence processing, database construction, and filtering criteria are provided in **Supplementary Methods S3**.

### Methanotroph and ammonia-oxidizing bacteria culture conditions

*M. trichosporium* OB3b and *M. album* BG8 were cultured in nitrate mineral salt (NMS) medium supplemented with 10 µM Cu under methane-containing headspace conditions, with 10% (v/v) methane in the headspace and the remaining volume filled with air^46^. *N. europaea* was grown under standard ammonia-oxidizing conditions^47^. Prior to experiments, methanotroph cultures were harvested during mid-exponential phase and adjusted to an optical density of OD_600_ = 0.3, whereas *N. europaea* cultures were adjusted to OD_600_ = 0.1 to account for differences in growth characteristics and cell density. Cultures were then used immediately for inhibition and kinetic assays.

### Methane oxidation kinetics and inhibition assays

Methane oxidation kinetics were measured in sealed 250 mL serum bottles containing actively growing cultures in NMS medium supplemented with 10 µM Cu. BNI compounds, including MHPP, 1,9-D, and LA, were added from stock solutions prepared in medium containing 0.01% (v/v) DMSO. Control treatments received equivalent DMSO without inhibitors. Final concentrations corresponded to estimated IC_50_ values for MHPP (730 µM) and 1,9-D (1076 µM), and IC_40_ for LA (57 µM). Serum bottles were sealed, amended with methane (6 mL), and incubated at 30 °C with shaking at 120 rpm. Headspace methane concentrations were quantified using gas chromatography equipped with a flame ionization detector (GC-FID), and methane oxidation rates were calculated from concentration changes over time. Methane oxidation kinetics were estimated from single-dose depletion experiments conducted at an initial methane concentration selected to capture the linear consumption range while minimizing biomass growth; under these conditions, OD_600_ increased by less than 10% (from 0.30 to ≤0.33), approximating initial-rate conditions for kinetic analysis. Kinetic parameters (V_max_ and K_m_) were estimated by fitting methane consumption rates to the Michaelis–Menten model under pseudo–initial rate conditions (OriginLab, Northampton, MA, USA).

### Reversibility and copper-dependent inhibition assays

To evaluate whether inhibition of methane oxidation by BNI compounds was reversible, *M. trichosporium* OB3b cultures were exposed to BNI compounds at their respective IC_50_ (MHPP and 1,9-D) or IC_40_ (LA) concentrations under the same conditions described above. Methane oxidation kinetics were first measured in triplicate cultures to quantify inhibited activity (V_max_). Following this measurement, triplicate cultures were pooled prior to centrifugation (5,000 × g, 10 min) to minimize variability associated with cell harvesting. Cells were washed once with fresh NMS medium to remove residual inhibitors, centrifuged again, and resuspended in fresh medium. The resuspended culture was then redistributed into triplicate serum bottles, amended with methane, and methane oxidation kinetics were re-measured as described above.

To assess whether inhibition was associated with pMMO, additional assays were conducted under copper-replete (+Cu) and copper-limited (–Cu) conditions to modulate expression of pMMO and sMMO, respectively. OB3b cultures used for copper-limited (–Cu) assays were preconditioned on Cu-free NMS agar plates prior to liquid cultivation to minimize residual copper carryover and facilitate sMMO expression^48^. Copper-replete conditions were established using NMS medium supplemented with 10 µM Cu, consistent with the conditions used in the kinetic assays, whereas copper-limited conditions were established using NMS medium without added Cu. Methane oxidation kinetics were then measured following exposure to BNI compounds at the same IC50 or IC40 concentrations as described above. V_max_ was measured for cultures treated with BNI compounds and corresponding controls (without BNI compounds) under both copper-replete (+Cu) and copper-limited (–Cu) conditions, and relative activity was calculated by normalizing V_max_ values of treated samples to their corresponding control within each condition.

### Structural modeling and molecular docking

The trimeric pMMO complex (PmoA–PmoB–PmoC) from *M. trichosporium* OB3b was modeled using both SWISS-MODEL^49^ and AlphaFold3^50^. Structural templates were selected based on sequence coverage and model quality metrics, including Global Model Quality Estimation (GMQE) and Quaternary Structure Quality Estimation (QSQE). The resulting models were evaluated by comparison with the crystal structure deposited in Protein Data Bank (PDB ID: 3CHX)^33^, and the model with the lowest root-mean-square deviation was selected for subsequent analyses. Molecular docking was performed to identify potential binding sites of BNI compounds. Docking simulations were conducted using SwissDock^51,52^ with the AutoDock Vina algorithm in blind docking mode^53^, allowing exploration of the entire protein surface exposed to the periplasm. Ligand structures were provided as SMILES inputs. Binding poses were ranked based on predicted binding energy (kcal mol⁻¹), and clusters of low-energy conformations were used to identify candidate binding regions. Binding sites that were inaccessible from the protein exterior, including internal cavities and membrane-embedded regions, were excluded from further analysis. Docking results were visualized and analyzed using PyMOL^54^. In addition to native BNI compounds, structural analogues of 1,9-D, including dodecanoic acid and 1-decanol, were selected to probe structure–activity relationships and evaluated using the same methane oxidation inhibition assay described above, under identical experimental conditions. IC_50_ values were determined by testing a range of concentrations and fitting the resulting data to the Michaelis–Menten model.

### RNA sequencing and transcriptomic analysis

RNA-seq analysis was performed to assess transcriptional responses of *M. trichosporium* OB3b to BNI compounds. Cultures were grown under the conditions described above and exposed to MHPP and 1,9-D at their estimated IC_50_ concentrations and to LA at its IC_40_ concentration. Prior to RNA-seq analysis, methane consumption over time was monitored under control and BNI-treated conditions to identify representative sampling points. Based on these profiles, samples were collected at 10 h, corresponding to peak methane oxidation activity to capture active transcriptional states (see **Supplementary Methods S4** and **Supplementary Fig. S2**). Total RNA was extracted using the MagMAX™ Microbiome Ultra Nucleic Acid Isolation Kit (Applied Biosystems, USA) in conjunction with the KingFisher™ Duo Prime automated nucleic acid extraction system (Thermo Fisher Scientific, USA), followed by rRNA removal prior to library preparation. Sequencing libraries were prepared using a strand-specific protocol and sequenced on an Illumina platform (Novogene Co., Ltd., China). Raw reads were quality-filtered using fastp to remove adapter sequences, poly-N reads, and low-quality reads. Clean reads were mapped to the reference genome of *M. trichosporium* OB3b (Genbank accession number GCA_002752655.1) using Bowtie2 (v2.5.4). Gene expression levels were quantified using featureCounts (v2.0.6), and normalized expression levels were calculated as fragments per kilobase of transcript per million mapped reads (FPKM) (see **Supplementary Table S3**). Differential expression analysis was performed using DESeq2 (v1.42.0), and p-values were adjusted using the Benjamini–Hochberg method. Genes with an adjusted p-value < 0.05 were considered significantly differentially expressed. Functional enrichment analysis was conducted using the clusterProfiler package (v4.8.1) for Gene Ontology (GO) terms. Transcriptomic data were visualized using PCA, volcano plots, and heatmaps. All treatments were performed in biological triplicates (n = 3). Additional details are provided in the **Supplementary Methods S4**.

### Comparative sensitivity and cell-normalized inhibition analysis

To compare the inhibitory effects of BNI compounds across methane-oxidizing and ammonia-oxidizing bacteria, additional inhibition assays were performed with *M. album* BG8 and *N. europaea* using the same general experimental framework described above. BNI compounds, including MHPP, 1,9-D, and LA, were tested across a range of concentrations to determine IC_50_ values. For methanotrophs (OB3b and BG8), inhibitory effects were quantified based on methane oxidation kinetics as described above. In contrast, for *N. europaea*, ammonia oxidation activity was assessed by measuring nitrite accumulation after 1 h of incubation, following conventional protocols used in BNI studies,^9,18,19^ with nitrite quantified by the Griess method. To account for differences in biomass across assays, inhibitory potency was normalized to cell abundance based on 16S rRNA gene copy numbers (see **Supplementary Methods S5** and **Supplementary Table S4**). Genomic DNA was extracted from cultures, and quantitative PCR (qPCR) targeting the bacterial 16S rRNA gene was performed using the CFX Opus 96 Real-Time PCR System (Bio-Rad, Hercules, CA, USA) in 20 μL reaction volumes containing SsoAdvanced™ Universal SYBR Green Supermix (Bio-Rad), 300 nM of each primer, and 2 μL of template DNA. The universal primer pair 1055F and 1392R was used for amplification. Gene copy numbers were converted to estimated cell abundance based on known 16S rRNA gene copy numbers per genome (one copy for OB3b^55^ and *N. europaea*^47^ and two copies for BG8^56^). Cell-normalized inhibitory potency was calculated by dividing IC_50_ values by the estimated cell abundance in each assay, yielding units of μM cell^−1^. For comparative visualization, cell-normalized IC_50_ values were log-transformed and expressed as −log_10_(IC_50_) to facilitate comparison across orders of magnitude. For LA, IC_50_ values could not be reliably determined within the tested concentration range; therefore, maximum observed inhibition was used for comparative analysis.

### Statistical Analysis

All statistical analyses were performed using R (v5.3) and GraphPad Prism (v10.0, GraphPad Software, USA). Data are presented as mean ± standard deviation (s.d.) unless otherwise indicated. Comparisons among multiple groups were evaluated using one-way analysis of variance (ANOVA) followed by Tukey’s post hoc test. Differences between paired conditions were assessed using two-tailed paired t-tests, and comparisons between two independent groups were performed using two-tailed unpaired t-tests. Correlations between calculated binding affinity and inhibitory potency were evaluated using Pearson’s correlation coefficient and Spearman’s rank correlation. Statistical significance was defined at P < 0.05. Where appropriate, more stringent significance levels (P < 0.01 and P < 0.001) were additionally reported.

## Acknowledgment

This work was supported by the National Science Foundation awards (no. 2144189 and 2435179).

