## Supplementary Information for "Biological nitrification inhibition compromises the soil methane sink"

This file contains:
– Supplementary Methods S1–S5
– Supplementary Figures S1–S2
– Supplementary Tables S1–S4

### Supplementary Methods

#### Supplementary Methods S1 | Soil characterization and column experiment.

The soil used in this study was classified as a silty clay loam according to the United States Department of Agriculture (USDA) soil taxonomy, consisting of 14% sand, 58% silt and 28% clay. The soil exhibited the following physicochemical properties: pH 7.3; total carbon (TC) 1.43%; total nitrogen (TN) 0.13%; total phosphorus (TP) 390 ppm; $NO_3^-$-N 8 ppm; $NH_4^+$-N 7 ppm; Ca 3111 ppm; Cu 1 ppm; Mg 131 ppm; Mn 9 ppm; Na 13 ppm; K 258 ppm; Zn 0.3 ppm; and Fe 29 ppm.

Acrylic soil columns (10 in internal diameter, 22 in length; **Fig. 1a**) were constructed from transparent tubing and fitted with airtight lids. Each column base was packed with a 2-cm layer of 3-mm glass beads to ensure uniform gas distribution. Soil was packed to a height of 25 cm, resulting in a bulk density of 1.9–2.0 g $cm^{-3}$. A gas mixture of $CH_4$ and $CO_2$ (50:50 v/v) was continuously supplied from the base of each column at a flow rate of 5 ± 0.5 mL $min^{-1}$, while humidified ambient air was introduced into the headspace at 20 mL $min^{-1}$ to simulate diffusive exchange. Soil moisture was monitored using an EC-5 probe (Decagon Devices, Pullman, WA, USA) and maintained at 45–55% water-holding capacity using an automated irrigation system. Ambient temperature and headspace relative humidity were monitored using a wireless thermo-hygrometer (Model H5074001, Govee, Hong Kong). Lighting was provided by LED strips mounted beneath the reactor lids, delivering approximately 200 μmol $m^{-2}$ $s^{-1}$ photosynthetically active radiation under a 16 h light / 8 h dark cycle. Experimental treatments (n = 3 per condition) included: (i) control soil; (ii) $NH_4^+$ amendment (25 mg $NH_4^+$-N·$kg^{-1}$ soil); and (iii) $NH_4^+$ amendment with *Brachiaria* seedlings Seedlings were transplanted at the 3–4 leaf stage. After two months of incubation, soil samples were collected from the rhizosphere (10–15 cm depth), homogenized, and processed for downstream analyses.

Methane oxidation kinetics were measured within 24 h of sampling. Soil subsamples (30 g, wet weight) were incubated in 160 mL serum bottles amended with 1% $CH_4$ (1.6 mL headspace). Bottles were incubated at 30 °C in the dark. Headspace samples (100 μL) were collected at 40-min intervals using gas-tight syringes and analyzed on an Agilent 7890 GC-FID. Methane oxidation rates were calculated from depletion curves, and kinetic parameters ($V_{max}$, $K_m$) were estimated by fitting the data to the Michaelis–Menten model using OriginLab (OriginLab Corporation, Northampton, MA, USA).

#### Supplementary Methods S2 | Methane oxidation kinetics and dose–response modeling.

Methane oxidation kinetics were measured in sealed 250 mL serum bottles containing actively growing cultures in nitrate mineral salts (NMS) medium supplemented with 10 μM Cu. BNI compounds (MHPP, 1,9-decanediol (1,9-D), and linoleic acid (LA)) were added from stock solutions prepared in medium containing 0.01% (v/v) DMSO. Control treatments received equivalent DMSO without BNI compounds. Serum bottles were sealed, amended with methane (6 mL), and incubated at 30 °C with shaking at 120 rpm. Headspace methane concentrations were monitored over time using gas chromatography equipped with a flame ionization detector (GC-FID), and methane oxidation rates were calculated from the linear decrease in methane concentration. Kinetic measurements were conducted under pseudo–initial rate conditions using single-dose depletion experiments. The initial methane concentration was selected to ensure linear consumption while minimizing biomass growth; under these conditions, $OD_{600}$ increased by less than 10% (0.30 to < 0.33), allowing approximation of initial-rate kinetics. These conditions are commonly used to approximate initial-rate kinetics, where substrate depletion and biomass growth do not significantly influence reaction rates. Methane oxidation kinetics were described using the Michaelis–Menten model:

$$v = \frac{V_{\max}[S]}{K_m + [S]}$$

where $v$ is the methane oxidation rate, $V_{max}$ is the maximum reaction rate, $K_m$ is the half-saturation constant, and [S] is the methane concentration in the headspace. Kinetic parameters ($V_{max}$ and $K_m$) were estimated by nonlinear regression using OriginLab (OriginLab Corporation, Northampton, MA, USA).

To quantify inhibition by BNI compounds, relative inhibition (%) was calculated based on the reduction in $V_{max}$ compared to untreated controls:

$$\text{Inhibition (\%)} = \left(1 - \frac{V_{\max\ ,\ treated}}{V_{\max\ ,\ control}}\right) \times 100$$

Dose–response relationships between inhibitor concentration and percent inhibition were fitted using a four-parameter logistic (4PL) model:

$$y = A_2 + \frac{A_1 - A_2}{1 + \left(\frac{x}{IC_{50}}\right)^p}$$

where $y$ is the observed inhibition (%), $x$ is the inhibitor concentration, $A_1$ and $A_2$ represent the minimum and maximum asymptotes, respectively, $IC_{50}$ is the inhibitor concentration corresponding to 50% inhibition, and $p$ is the Hill slope describing the steepness of the curve. Parameters were estimated by nonlinear least-squares regression in OriginLab. For LA, inhibition did not reach 50% within the tested concentration range; therefore, $IC_{40}$ values were estimated from the fitted 4PL model and used for subsequent analyses.

**Supplementary Methods S3 | Metagenomic analysis of methanotroph communities.**
Metagenomic sequencing was performed to characterize methanotroph community composition in soil bioreactor samples. Genomic DNA was extracted using the MagMAX™ Microbiome Ultra Nucleic Acid Isolation Kit (Applied Biosystems, USA) with the KingFisher™ Duo Prime system. Libraries were prepared and sequenced on an Illumina platform (Novogene Co., Ltd., China). Paired-end reads were merged using FLASH (parameters: -m 10, -M 65, -x 0.25) and converted to FASTA format using Seqtk. High-quality sequences were used for downstream analysis. Custom *pmoA* reference databases were constructed using curated methanotroph sequences (**Supplementary Table S2**) and supplemented with selected FunGene entries to improve taxonomic resolution. Reference sequences were categorized into Type I, Type II, and

Type III methanotrophs, and non-overlapping databases were generated for each group. Metagenomic sequences were aligned against the custom *pmoA* databases using BLASTN with an e-value threshold of $1 \times 10^{-6}$ and a minimum identity of 80%. The highest-scoring hit for each query was retained for classification. Relative abundances of *pmoA*-containing methanotroph lineages were compared across treatments.

**Supplementary Methods S4 | RNA sequencing and transcriptomic analysis.**
RNA-seq analysis was performed to assess transcriptional responses of *Methylosinus trichosporium* OB3b to BNI compounds. Cultures were exposed to MHPP and 1,9-D at their estimated $IC_{50}$ concentrations and to LA at its $IC_{40}$ concentration. Samples were collected at 10 h, corresponding to a period of high methane oxidation activity as determined from preliminary time-course experiments. Total RNA was extracted using the MagMAX™ Microbiome Ultra Nucleic Acid Isolation Kit (Applied Biosystems, USA) in conjunction with the KingFisher™ Duo Prime automated nucleic acid extraction system (Thermo Fisher Scientific, USA). Ribosomal RNA was removed prior to library preparation. Strand-specific libraries were prepared and sequenced on an Illumina platform (Novogene Co., Ltd., China). Raw reads were quality-filtered using fastp to remove adapter sequences, poly-N reads, and low-quality reads. Clean reads were mapped to the reference genome of *M. trichosporium* OB3b (GenBank accession number GCA_002752655.1)[1] using Bowtie2 (v2.5.4). Gene expression levels were quantified using featureCounts (v2.0.6) and normalized as fragments per kilobase of transcript per million mapped reads (FPKM). Differential expression analysis was performed using DESeq2 (v1.42.0), with *p*-values adjusted using the Benjamini–Hochberg method. Genes with an adjusted p-value $< 0.05$ were considered significantly differentially expressed. Functional enrichment analysis was conducted using clusterProfiler (v4.8.1) for Gene Ontology (GO) terms.

**Supplementary Methods S5 | Quantitative PCR analysis.**
Genomic DNA contamination in RNA samples was assessed by PCR using universal 16S rRNA gene primers on DNase-treated RNA prior to reverse transcription. Total RNA was reverse-transcribed to cDNA using Invitrogen SuperScript™ IV Reverse Transcriptase (Thermo Fisher Scientific, USA) according to the manufacturer's instructions. Transcript abundance was quantified by RT–qPCR using a CFX Opus 96 Real-Time PCR System (Bio-Rad, Hercules, CA, USA). Reactions were performed in 20 μL containing 10 μL of SsoAdvanced™ Universal SYBR Green Supermix (Bio-Rad), 300 nM of each primer, and 2 μL of template cDNA. Quantification was performed using standard curves generated from serial 10-fold dilutions of cloned plasmids or double-stranded synthetic DNA fragments (gBlocks®, Integrated DNA Technologies, Coralville, IA, USA), with each standard measured in triplicate. Relative *pmoA* transcript abundance was normalized to 16S rRNA gene copy number.

### Supplementary Figures

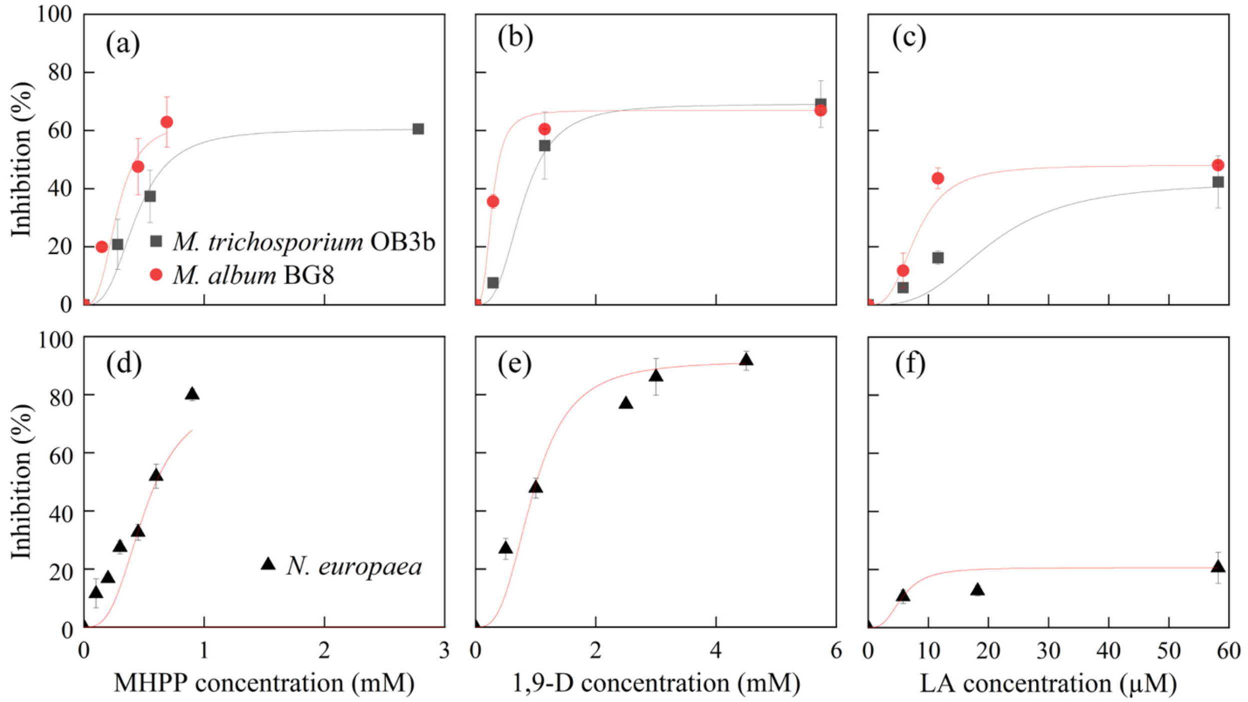


**Figure S1 | Concentration-dependent inhibition of methane- and ammonia-oxidizing bacteria by BNI compounds.**

(a–c) Inhibition of methane oxidation in *Methylosinus trichosporium* OB3b (Type II; squares) and *Methylomicrobium album* BG8 (Type I) as a function of increasing concentrations of (a) methyl 3-(4-hydroxyphenyl)propionate (MHPP), (b) 1,9-decanediol (1,9-D), and (c) linoleic acid (LA).

(d–f) Inhibition of ammonia oxidation in *Nitrosomonas europaea* as a function of increasing concentrations of (d) MHPP, (e) 1,9-D, and (f) LA.

Inhibition (%) was calculated relative to untreated controls based on methane or ammonia oxidation activity. Solid lines represent fitted dose–response curves using a four-parameter logistic (4PL) model. Data points represent mean ± s.d. (n = 3).

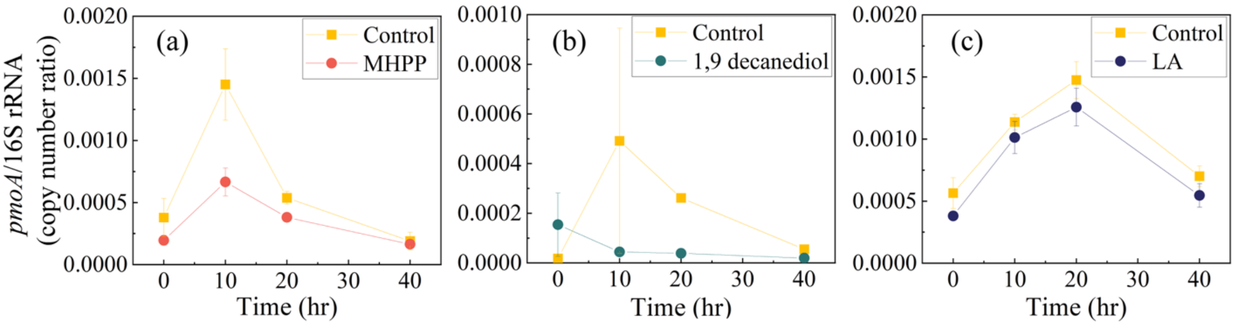


**Figure S2 | Time-dependent changes in *pmoA* transcript abundance in response to BNI compounds.**

Relative *pmoA* transcript levels (normalized to 16S rRNA gene copy number) were monitored over time following exposure to MHPP, 1,9-D, and LA. Across treatments, *pmoA* transcript levels exhibited dynamic temporal responses, with 10 h representing a time point capturing strong or near-peak transcriptional activity depending on the treatment, supporting the selection of this time point for RNA-seq analysis.

### Supplementary Tables

**Table S1 | Summary of fitted inhibitory concentrations and maximal inhibition values for BNI compounds across methane- and ammonia-oxidizing bacteria.**

IC50 values were obtained from four-parameter logistic fits for MHPP and 1,9-D. Because inhibition by LA did not reach 50% within the tested concentration range, $IC_{40}$ values are reported for OB3b and BG8, whereas maximum observed inhibition is reported for *N. europaea*. OB3b and BG8 refer to *Methylosinus trichosporium* OB3b and *Methylomicrobium album* BG8, respectively.

| Compound | Organism | Metric | Value | Unit | Notes |
|---|---|---|---|---|---|
| MHPP | OB3b | $IC_{50}$ | 0.73 | mM | |
| | BG8 | $IC_{50}$ | 0.43 | mM | |
| | *N. europaea* | $IC_{50}$ | 0.60 | mM | |
| 1,9-D | OB3b | $IC_{50}$ | 1.07 | mM | |
| | BG8 | $IC_{50}$ | 0.39 | mM | |
| | *N. europaea* | $IC_{50}$ | 1.00 | mM | |
| LA | OB3b | $IC_{40}$ | 53 | µM | $IC_{50}$ not reached |
| | BG8 | $IC_{40}$ | 14 | µM | $IC_{50}$ not reached |
| | *N. europaea* | max inhibition | 13 | % | at 57 µM; $IC_{50}$ not reached |

**Table S2 | Curated methanotroph *pmoA* reference sequences used for database construction.**

Reference sequences were compiled through a comprehensive literature survey of all characterized methanotroph species reported as of December 2025. Candidatus taxa were excluded to ensure inclusion of taxonomically validated species only. For entries derived from whole-genome or contig accessions, *pmoA* sequences were identified from annotated genomic regions. Because *pmoA* and *amoA* belong to the homologous copper membrane monooxygenase (CuMMO) family, some *pmoA* genes may be annotated as *amoA* in public databases. These sequences were manually verified and curated based on sequence similarity and taxonomic context prior to inclusion.

| Species | Methanotrophs type | GenBank accession |
|---|---|---|
| *Methylomonas albis* | Type I | KX958422 |
| *Methylomonas defluvii* | Type I | OR881399 |
| *Methylomonas fluvii* | Type I | KX958424 |
| *Methylomonas koyamae* | Type I | OR004531 |
| *Methylomonas lenta* | Type I | HF954359 |
| *Methylomonas methanica* | Type I | EU722434 |
| *Methylomonas paludis* | Type I | HE801217 |
| *Methylomonas rapida* | Type I | MK165450 |
| *Methylomonas rivi* | Type I | GCA_024505165.1 |
| *Methylomonas rosea* | Type I | GCA_024505055.1 |
| *Methylomonas subterranea* | Type I | GCA_024505185.1 |
| *Methylovulum miyakonense* | Type I | AB501285 |
| *Methylovulum psychrotolerans* | Type I | KT381579 |
| *Methylomicrobium agile* | Type I | GCA_000733855.1 |
| *Methylomicrobium album* | Type I | FJ713039 |
| *Methylomicrobium alcaliphilum* | Type I | NC_016112 |
| *Methylomicrobium buryatense* | Type I | AF307139 |
| *Methylomicrobium kenyense* | Type I | JN687579 |
| *Methylomicrobium japanense* | Type I | AB253367 |
| *Methylomicrobium pelagicum* | Type I | U31652 |
| *Methylocucumis oryzae* | Type I | MT366581 |
| *Methylobacter marinus* | Type I | KB912877 |
| *Methylobacter luteus* | Type I | LC744973 |
| *Methylobacter tundripaludum* | Type I | AJ414658 |
| *Methylobacter psychrophilus* | Type I | OK157433 |
| *Methyloprofundus sedimenti* | Type I | KF484908 |
| *Methylosoma difficile* | Type I | DQ119047 |
| *Methyloglobulus morosus* | Type I | JN386975 |
| *Methylosarcina fibrata* | Type I | AF177325 |
| *Methylosarcina quisquiliarum* | Type I | AF177326 |
| *Methylococcus capsulatus* | Type I | AF533666 |
| *Methylococcus geothermalis* | Type I | MN735154 |
| *Methylococcus mesophilus* | Type I | CP110921 |
| *Methylocaldum gracile* | Type I | OQ354220 |
| *Methylocaldum marinum* | Type I | AB900159 |
| *Methylocaldum szegediense* | Type I | U89303 |
| *Methylocaldum tepidum* | Type I | U89304 |
| *Methylotetracoccus oryzae* | Type I | GCA_006175985.1 |
| *Methylomagnum ishizawai* | Type I | AB669168 |
| *Methyloparacoccus murrellii* | Type I | AB636304 |
| *Methylogaea oryzae* | Type I | EU359002 |
| *Methylospira mobilis* | Type I | KU216207 |
| *Methyloterricola oryzae* | Type I | KY129803 |

| | | |
|---|---|---|
| *Methylohalobius crimeensis* | Type I | AJ581836 |
| *Methylothermus subterraneus* | Type I | AB536748 |
| *Methylomarinovum caldicuralii* | Type I | AB302948 |
| *Crenothrix polyspora* | Type I | DQ295904 |
| *Clonothrix fusca* | Type I | DQ984192 |
| *Methylocapsa acidiphila* | Type II | AJ278727 |
| *Methylocapsa aurea* | Type II | FN433470 |
| *Methylocapsa palsarum* | Type II | KP715290 |
| *Methylocapsa polymorpha* | Type II | CP136862 |
| *Methylosinus sporium* | Type II | FJ713041 |
| *Methylosinus trichosporium* | Type II | AJ868409 |
| *Methylocystis bryophila* | Type II | FN422005 |
| *Methylocystis echinoides* | Type II | AJ459000 |
| *Methylocystis heyeri* | Type II | AM283546 |
| *Methylocystis hirsuta* | Type II | DQ364434 |
| *Methylocystis iwaonis* | Type II | AB669161 |
| *Methylocystis parva* | Type II | MF614620 |
| *Methylocystis rosea* | Type II | AJ414657 |
| *Methylocystis silviterrae* | Type II | GCA_013350005.1 |
| *Methylocystis suflitae* | Type II | GCA_024448135.1 |
| *Methylacidimicrobium tartarophylax* | Type III | GCA_902143375.2 |
| *Methylacidimicrobium cyclopophantes* | Type III | GCA_902143385.2 |
| *Methylacidimicrobium thermophilum* | Type III | LR797830 |
| *Methylacidiphilum infernorum* | Type III | CP000975 |
| *Methylacidiphilum fumariolicum* | Type III | EF591085 |
| *Methylacidiphilum kamchatkense* | Type III | CP037899 |
| *Methylacidiphilum caldifontis* | Type III | CP065957 |

For entries with whole-genome or contig accessions, *pmoA* sequences were identified from annotated genomic regions and extracted prior to database construction.

**Table S3 | Summary of RNA-seq quality and mapping statistics across treatments and biological replicates.**

| Treatment | Replicate | Clean reads | Mapping rate (%) | Unique mapping (%) | Q30 (%) | GC (%) |
|---|---|---|---|---|---|---|
| Control | 1 | 28769118 | 98.52 | 81.90 | 94.99 | 64.51 |
| | 2 | 28262112 | 98.38 | 82.36 | 94.84 | 65.04 |
| | 3 | 26919952 | 98.41 | 83.24 | 95.10 | 64.52 |
| MHPP | 1 | 23990292 | 98.57 | 80.13 | 94.94 | 64.74 |
| | 2 | 24810868 | 98.54 | 82.53 | 94.88 | 65.11 |
| | 3 | 26074358 | 98.44 | 83.30 | 94.86 | 64.75 |
| 1,9-D | 1 | 26912524 | 97.12 | 81.46 | 94.86 | 64.72 |
| | 2 | 26312280 | 98.24 | 82.31 | 94.94 | 64.71 |
| | 3 | 25350700 | 98.18 | 83.47 | 94.93 | 64.90 |
| LA | 1 | 25372204 | 97.47 | 81.28 | 95.01 | 64.78 |
| | 2 | 28247442 | 95.05 | 77.59 | 95.12 | 64.60 |
| | 3 | 31439004 | 97.10 | 80.01 | 94.90 | 64.64 |

**Table S4 | Primers used for quantitative PCR and detection ranges.**

| Target gene | Primer | Primer sequences (5'-3') | Quantification range | References |
|---|---|---|---|---|
| 16S rRNA | 1055F<br>1392R | ATGGCTGTOGTCAGCT<br>ACGGGCGGTGTGTAC | $10^2$-$10^9$ | [2] |
| *pmoA* | A189<br>mb661 | GGNGACTGGGACTTCTGG<br>CCGGMGCAACGTCYTTACC | $10^2$-$10^9$ | [3] |

Abbreviations for degenerate nucleotide positions are as follows: N = A, C, G, or T; M = A or C; Y = C or T.

References
